# Biases in population codes with a few active neurons

**DOI:** 10.1101/2024.09.20.614069

**Authors:** Sander Keemink, Mark C.W. van Rossum

**Affiliations:** Department of Machine Learning and Neural Computing, Donders Institute for Brain, Cognition and Behaviour, Radboud University, Nijmegen 6525 GD, NL; School of Psychology and School of Mathematical Sciences, University of Nottingham, Nottingham NG7 2RD, UK

## Abstract

Throughout the brain information is coded in the activity of multiple neurons at once, so called population codes. Population codes are a robust and accurate way of coding information. One can evaluate the quality of population coding by trying to read out the code with a decoder, and estimate the encoded stimulus. Coding quality has traditionally been evaluated in terms of the trial-to-trial variation in the estimate. However, codes can also display biases. While most decoders yield unbiased estimators in the limit of many active neurons, we find that when only few neurons are active, biases readily emerge for many decoders. We show that the biases turn out to have a non-trivial dependence on noise and tuning curve properties. We also introduce a technique to estimate the bias and variance of Bayesian decoders. Overall, the work expands our understanding of population coding.

---

In many brain areas neurons code information in the form of a population code. That is, a stimulus drives the activity of many neurons, thereby yielding robustness to neural noise and neural death, while still having a high capacity (Pouget et al., 2000). As population codes are at the core of neural information processing, over the years many properties of population codes have been studied. The central theme of many of these studies has been the quality of the code. One can quantify the code quality by the trial-to-trial variance in the decoded variable. Various studies have addressed questions, such as, how accurate are population codes in the presence of noise (Paradiso, 1988), how do correlations in the noise degrade information (Abbott and Dayan, 1999; Wu et al., 2002), and how can population codes be transmitted with minimal accuracy loss (Renart and van Rossum, 2012).

In addition to trial-to-trial variance, a decoder can also exhibit a bias. That is, even after averaging over many trials a systematic difference between the true value of the encoded stimulus and its estimate remains. Given an estimate 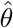 and a true stimulus value Θ, one defines bias as

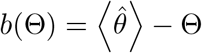

where the angular brackets denote averaging over trials. In the limit of many active neurons with low noise and with perfect knowledge of the encoding system, commonly used decoders have no bias. But biases can emerge when these assumptions are not met. For instance, biases emerge when the encoder adapts but the decoder does not, so that the decoder model does not match the encoder (Seriès et al., 2009). Another case in which biases emerge is when multiple stimuli are coded simultaneously in a neural population (Amari and Burnashev, 2003; Keemink et al., 2018a). Biases have also been studied in Bayesian perception when the stimulus priors are non-uniform (Wei and Stocker, 2015). The purpose of this paper is to explore biases in sparse populations with uniform stimulus priors.

## Setup

When discussing population codes, one typically thinks of systems with many neurons, but in insects populations might be tiny. A common example of a low-dimensional population code is the cricket wind sensor system (Miller et al., 1991; Theunissen and Miller, 1991; Salinas and Abbott, 1994). Here an angular stimulus Θ, the wind direction, is encoded by just four neurons, *k* = 1 … *N*, with *N* = 4. We assume that the neurons have preferred stimuli *ϕ*_*k*_ that are spaced 90 degrees apart (*ϕ*_*k*_ = *kπ/*2) and have rectified cosine tuning

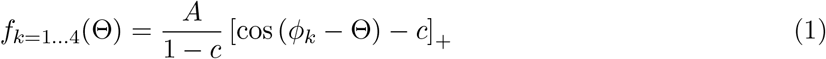

where *A* is the response amplitude. The threshold *c* determines the width of the tuning curves and hence their overlap; a large value leads to narrow tuning (−1 ≤ *c <* 1). We assume Gaussian additive, uncorrelated noise, so that on a given trial the response of neuron *i* is *r*_*k*_ = *f*_*k*_(Θ) + *σν*, where *σ* is the standard deviation of the noise, and *ν* is a sample from a Gaussian distribution with unit variance. We have also examined Poisson noise, and found it made no qualitative difference to our findings. We set amplitude *A* = 1, and unless denoted otherwise use parameters *c* = −0.1, *σ* = 0.1 (The cricket tuning curves were in Salinas and Abbott (1994) fitted with *c* = −0.14). The corresponding tuning curves are illustrated in Fig. 1A.

**Figure 1:**
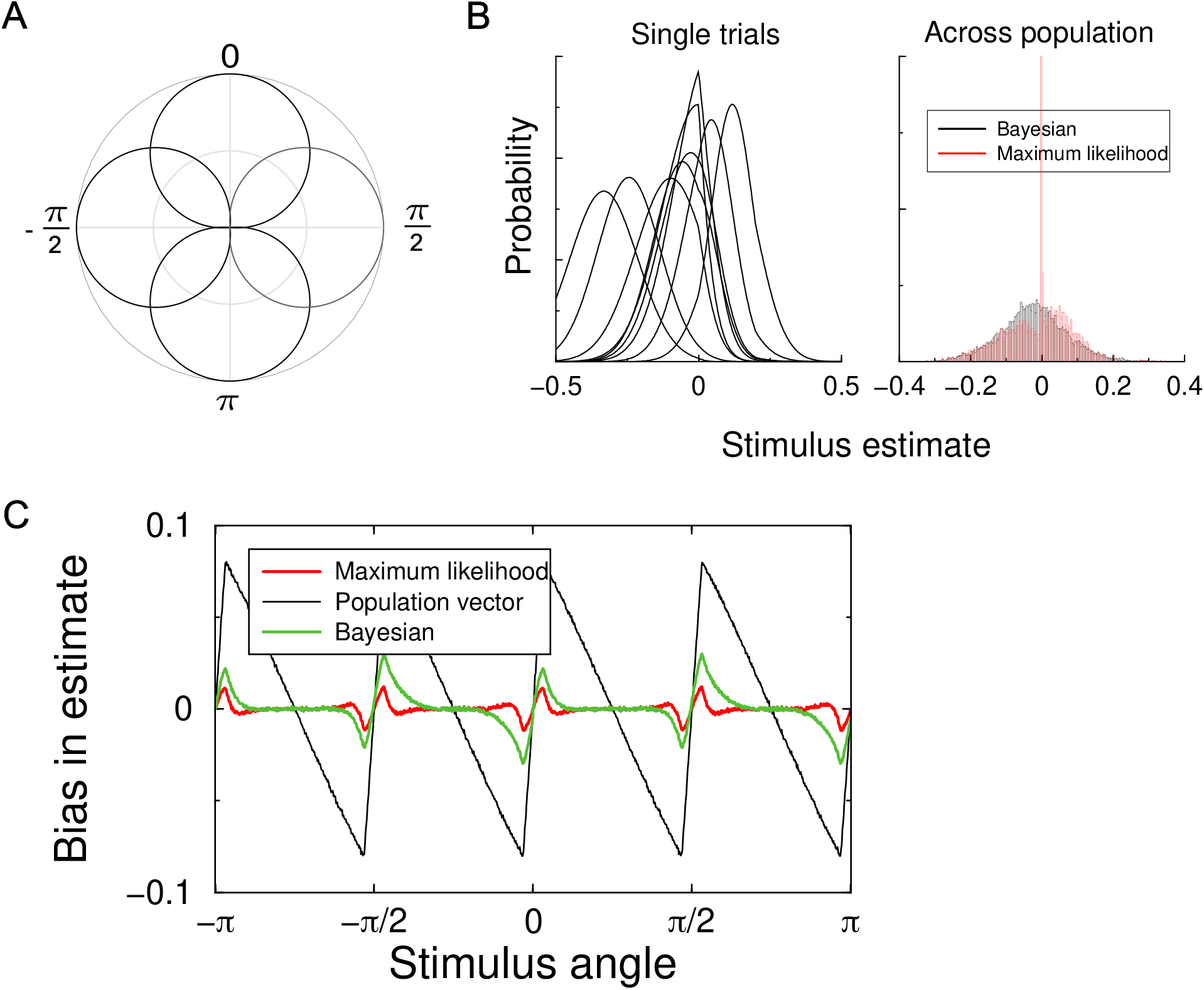
Estimator bias in a model of the four neuron cricket wind direction system. A. The tuning curves of the 4 neurons in a polar plot as a function of the encoded stimulus angle. The distance from the center corresponds to the firing rate. B. Left: The estimate distribution for a few trials. On a given trial, a Bayesian estimate extracts the mean of the distribution; the maximum likelihood uses the maximum. Right: the distribution of estimates; the bias was in this case -0.023 for the Bayesian and -0.012 for ML decoder. The distributions are plotted relative to true stimulus angle, which was -0.1. C. Estimator bias. All commonly used decoders show bias. The preferred angles correspond to the tick-marks. The bias is in this case largely repulsive, away from the preferred stimuli.

The task of a decoder is to estimate the true value of the encoded stimulus from the noisy response vector ***r***. We compare a number of commonly used decoders (Dayan and Abbott, 2001):

1. The *population vector* decoder: On a given trial one first constructs the population vector 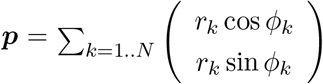. From ***p*** the angle is estimated as 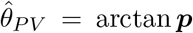.This decoder is identical to the maximum likelihood decoder in the case of dense von Mises tuning curves (see below) with Poisson noise (Snippe, 1996; Keemink et al., 2018b), but in general its trial-to-trial variance is larger than for the other estimators, that is, it is not efficient (see below).
2. The *maximum likelihood decoder* : According to Bayes’ theorem, the probability for a certain stimulus given the response is *P* (*θ*|***r***) = *P* (***r***|*θ*)*P* (*θ*)*/P* (***r***), Fig.1B left. Under the assumption of a uniform flat prior *P* (*θ*) and Gaussian noise, one has log 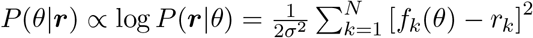.The maximum likelihood estimate is the stimulus angle that maximizes the likelihood, i.e. 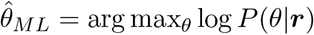. Numerically, the estimate can be found be minimizing the likelihood. In practice it is often easier to find the minimum in a finely spaced array of candidate stimuli.
3. The *B*ayesian decoder: The *Bayesian decoder* also relies on *P* (*θ*|***r***), but finds the estimate that minimizes a cost function. For the mean squared error cost, this is the mean of the distribution, hence 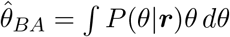.

### Emergence of bias

To examine the emergence of bias, we sample many noisy population responses and estimate the stimulus on each trial according to these three estimators, focussing on the stimulus angles close to one neuron’s preferred stimulus. In the limit where many, low noise neurons are active, the likelihood becomes a Gaussian, and as a result the maximum likelihood estimator equals the Bayes estimator when the prior is uniform. However, when there are just a few noisy neurons active this is no longer true. For instance, Figure 3.7 in the textbook by Dayan and Abbott (2001) shows a subtle difference in the variance of the ML and Bayesian decoders, hinting at a non-Gaussian posterior distribution. Also here the distribution of estimate is non-Gaussian, Fig.1B right.

**Figure 2:**
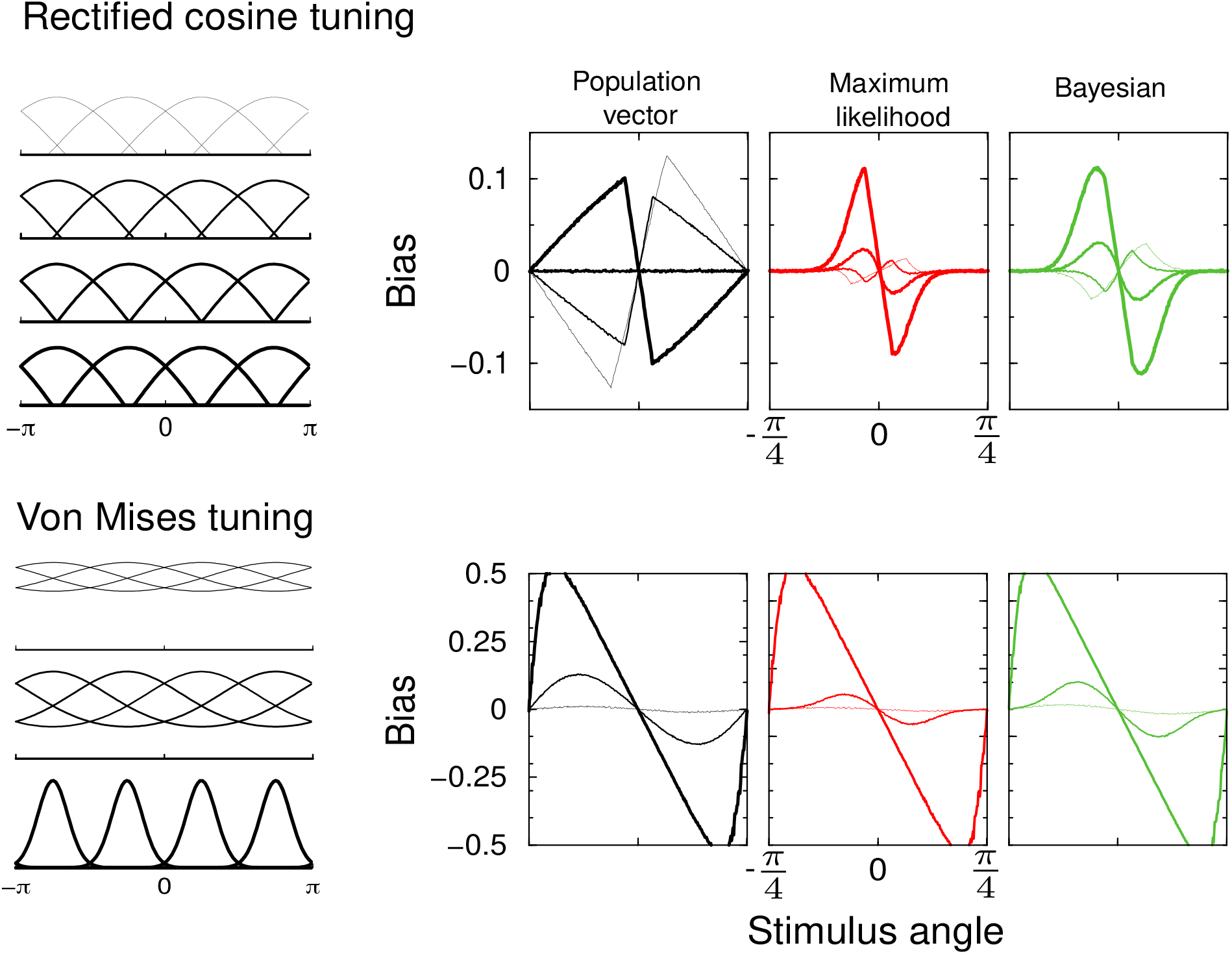
Top: Estimator bias for population vector (left), maximum likelihood (middle) and Bayesian (right) estimators for 4 different tuning curve widths (illustrated on the left). Rectified cosine tuning. Thin to thick curve *c* = −0.2, −0.1, 0, and 0.1. Gaussian noise *σ* = 0.1. Bottom: Estimator bias for von Mises tuning curves. In this case, bias is always attractive. Widths *w* = 2 (wide; thin curve), 0.5, and 0.1 (narrow; thickest curve).

**Figure 3:**
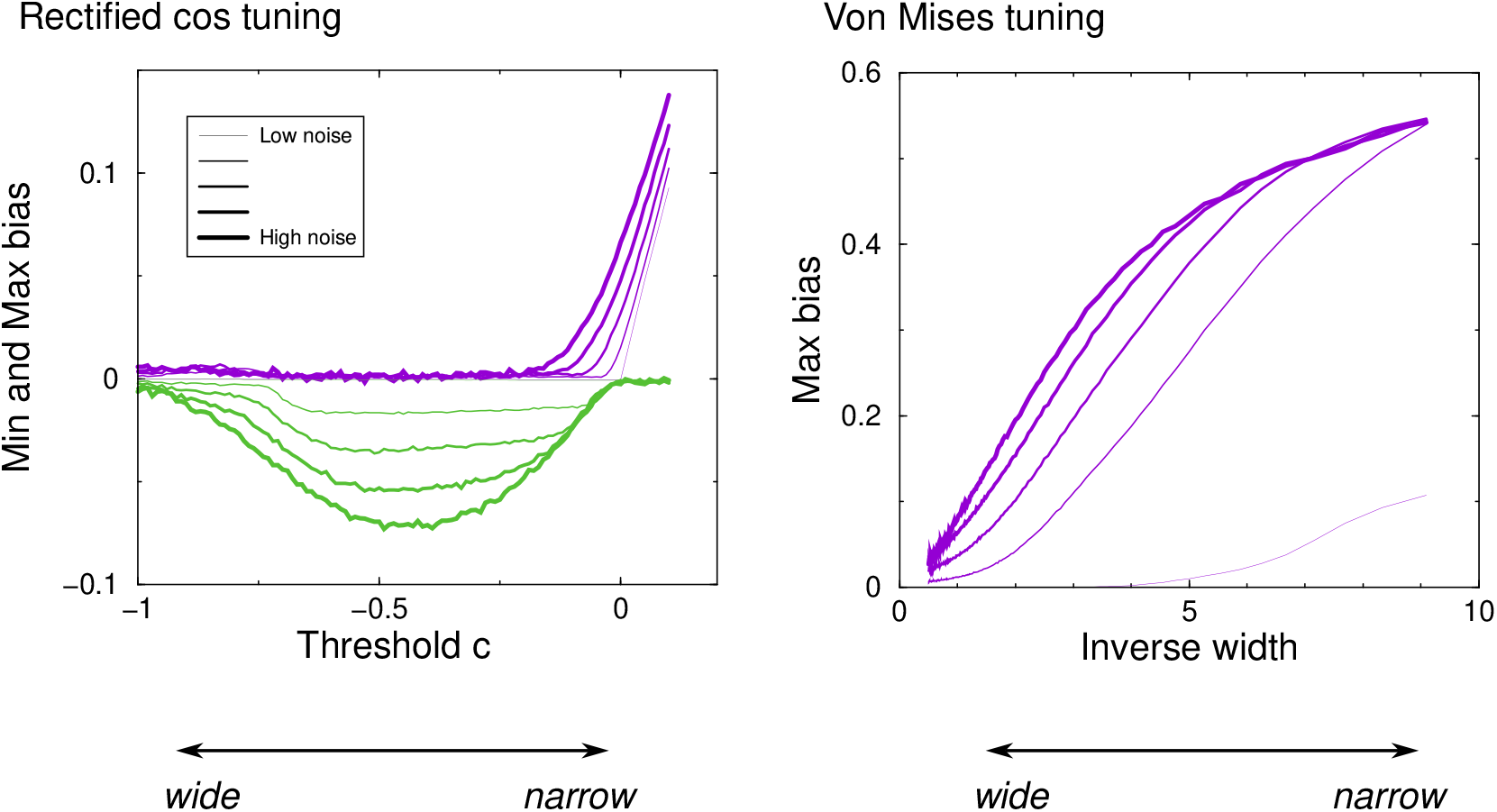
Left: Minimum (green) and maximum (purple) bias for the Bayesian decoder vs the tuning curve threshold. 5 different levels of Gaussian noise (*σ* = 0.01, 0.05, 0.1, 0.15, 0.2). Positive bias corresponds to attractive bias. Right: As left, but for von Mises tuning as a function of inverse width 1*/w*. In this case the bias is always positive.

The bias in the cricket wind direction system as a function of the stimulus angle w.r.t the preferred angle of one of the neurons is shown in Fig.1C. Due to symmetry, the bias is an odd, periodic function, zero at the preferred angles and halfway two preferred angles. All estimators display a bias that happens in this case to be mostly repulsive from the preferred direction: angles slightly less than the preferred angle, are estimated as being even less. The population vector has the largest bias, while the ML and Bayesian decoders have similar bias.

### Role of tuning curve shape and width

The tuning curve width is set by the threshold parameter *c* in Eq.1. To examine its role, we varied it and plot the bias, Fig.2. With wide tuning curves (thin curves), the bias is repulsive. As the tuning curves narrow, the bias becomes bi-phasic with both attractive and repulsive regions, or fully attractive, that is, stimuli near a preferred direction are estimated nearer to that direction. In the regime of narrow tuning (*c >* 0), stimuli close to the preferred angles activate just one neuron.

Consider a the neuron with preferred angle zero. When just this neuron is active, any off-peak response is ambiguous – the stimulus could have been on either side of the preferred stimulus, and both Θ, but also −Θ are equally likely. Noise will break this symmetry, hence estimating either stimulus equally. In other words, for *any* reasonable estimator the estimate will in such a case average to zero. That is, the bias is complete and attractive, *b*(Θ) = −Θ. The slope of the bias for all estimators is exactly -1 for angles close to zero (thick curves).

We also examine von Mises tuning curves, *f*_*k*=1…4_(Θ) ∝ exp [(cos(*ϕ*_*k*_ − Θ) − 1)*/w*]. In this case the bias is always attractive and increases for narrow tuning widths *w*, Fig. 2bottom. These results demonstrate that biases readily emerge in population codes with just a few neurons. While the three decoders always show similar biases, the dependence of the bias on tuning curve shape is intricate.

### Role of noise

In the above simulations the neural responses were noisy. Next, we investigate the effect of the strength of neural noise on the bias. We show results for the Bayesian decoder only, as the ML decoder is similar. To summarize the complicated bias curves, we extract the minimum and maximum bias of the Bayesian decoder across the range of stimulus angles on the left of the preferred stimulus (−*π/*4 ≤ Θ ≤ 0), so that a positive bias corresponds to attraction to the preferred stimulus. The minimum and maximum bias are plotted against the threshold parameter that sets the tuning width and for a number of noise levels, Fig.3 top. The noise level is indicated by the line thickness (thicker lines signify more noise). At the most negative threshold (*c* = −1), the tuning curves are pure cosines (broad tuning) and bias is minimal. For intermediate thresholds (*c* ≈ 0.5), the bias negative and at a given threshold, the curves in the figure are spaced out equally in the vertical direction, in other words the bias is proportional to the noise. In the region −0.2 ≲ *c* ≲ 0, both minimum and maximum bias are non-zero; here the bias profile is bi-phasic, as can be observed in Fig.1. For positive threshold (narrow tuning) we enter the ambiguous regime described above and bias persists even in the absence of noise.

For von Mises tuning, the bias is positive across all noise levels, Fig.3b. Again, while bias increases monotonically with noise, the dependence is non-linear (the curves are not equidistantly spaced vertically)

### Scaling with population size

The above results raise the question whether biases persist if the population contains more neurons. First we increased the number of neurons without changing any of the tuning curve properties, Fig. 4 (black curve). Next, we scaled the tuning such that the overall mean activity remained constant. First we decreased the amplitude as the number of neurons increased. When the amplitude was scaled, bias still reduced when using more neurons but less so (blue curve). In other words, the bias only depends weakly on the signal-to-noise ratio of the neurons, consistent with the above observations. As an alternative we scaled the tuning curve width, so that the average number of active neurons remains the same (red curve). In this case the bias dropped rapidly as well. In the limit of many narrowly tuned neurons, the bias scales as 1*/N*, that is, it is proportional to the distance between neurons. Such behavior is consistent with a generic scaling argument.

**Figure 4:**
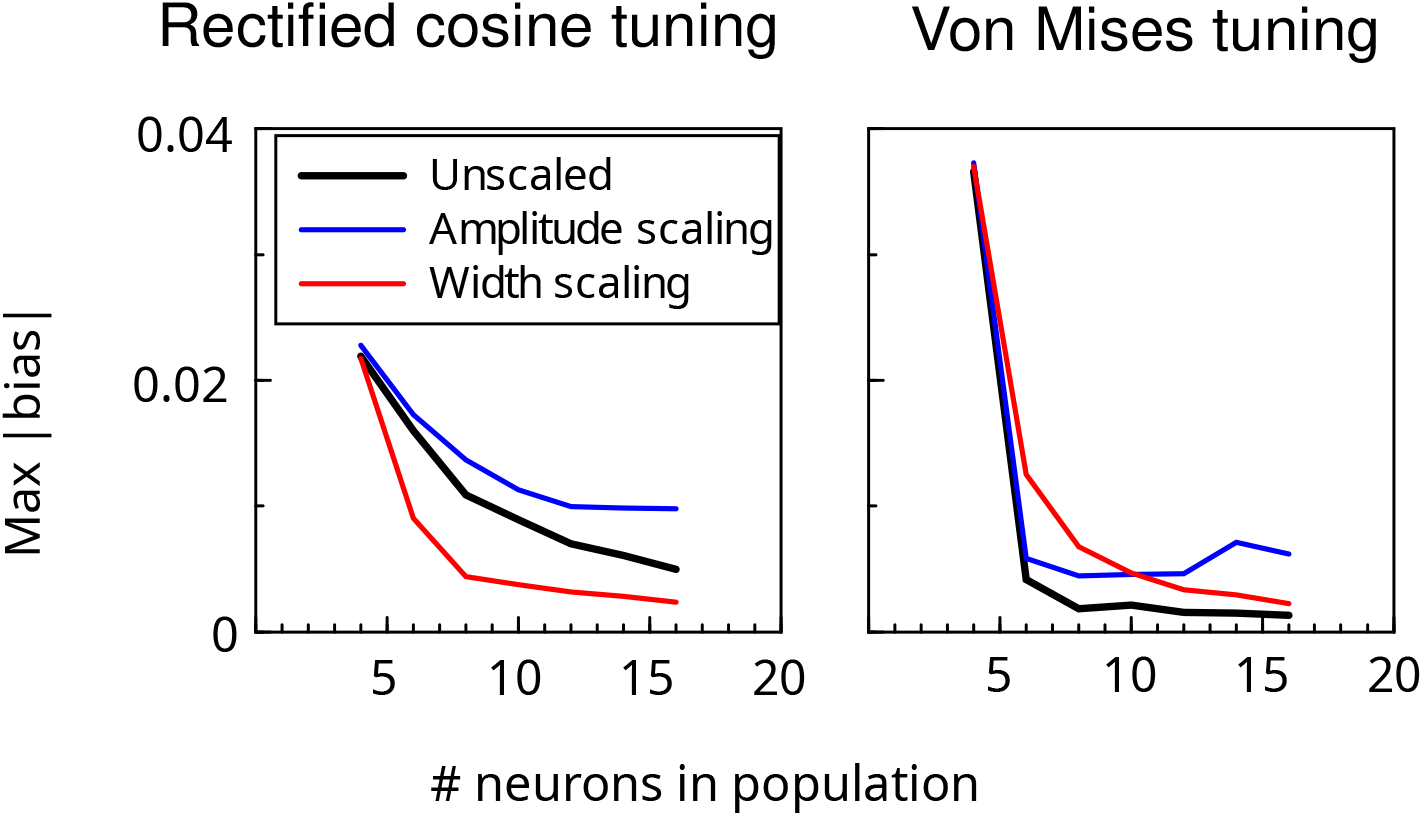
Maximum absolute bias vs the number of neurons in the population for the Bayesian decoder. Bias decreases when there are more neurons in the population. With amplitude scaling the mean population activity was kept the same by scaling down the response amplitude for larger population (blue curve). With width scaling this was achieving by narrowing the tuning curves. Threshold parameter *c* = −0.1 for the rectified cosine tuning with 4 neurons, and width *w* was 1 for von Mises tuning.

For von Mises tuning, Fig. 4 right, there is a much steeper decrease for all scaling methods. While it would be interesting to examine the scaling of the bias at larger numbers of neurons, this is numerically challenging. The reason is that as the bias diminishes, it becomes comparable to the trial-to-trial fluctuations in the estimates. To prevent this, an unworkable large number of simulated trials would be needed.

### Decoding efficiency

Finally, we study the interaction between the bias and decoder variance. The variance 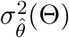 that any decoder can achieve is lower limited by the Cramer-Rao bound. For unbiased estimators it has a lower bound,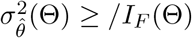, where for Gaussian noise the Fisher Information 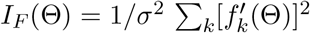. It is known that only for some estimation problems unbiased, minimal variance estimators exist (Kay, 1993). Indeed, when the noise is large the variance is larger than the bound (Xie, 2002). Nevertheless in the limit of many neurons and low noise, well known decoders such as maximum likelihood decoders reach the lower bound.

When the decoder has a bias *b*(Θ), the bound is

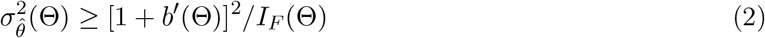

where *b*^*′*^(Θ) is the derivative of the bias w.r.t. the stimulus. One defines efficiency of the estimator as the ratio of right and left hand sides of Eq.2.

Due to the limited number of neurons, the system is not homogeneous and the variance of the decoder varies across with the stimulus, Fig. 5 (green curve). It drops near the preferred stimulus, as there three instead of two neurons are simultaneously active, making a more accurate estimate possible. But it increases again near zero as there the most active neuron has zero slope and won’t contribute to the estimate. The bias corrected Cramer-Rao bound drops as well, so that the Cramer-Rao bound holds (green curve lies above black curve). In fact, while the efficiency is close to one away from the preferred stimulus (green and black curves are close), it drops below one near the preferred stimulus. Note, the Cramer–Rao bound shows substantial numerical noise as it relies on the derivative of the bias.

**Figure 5:**
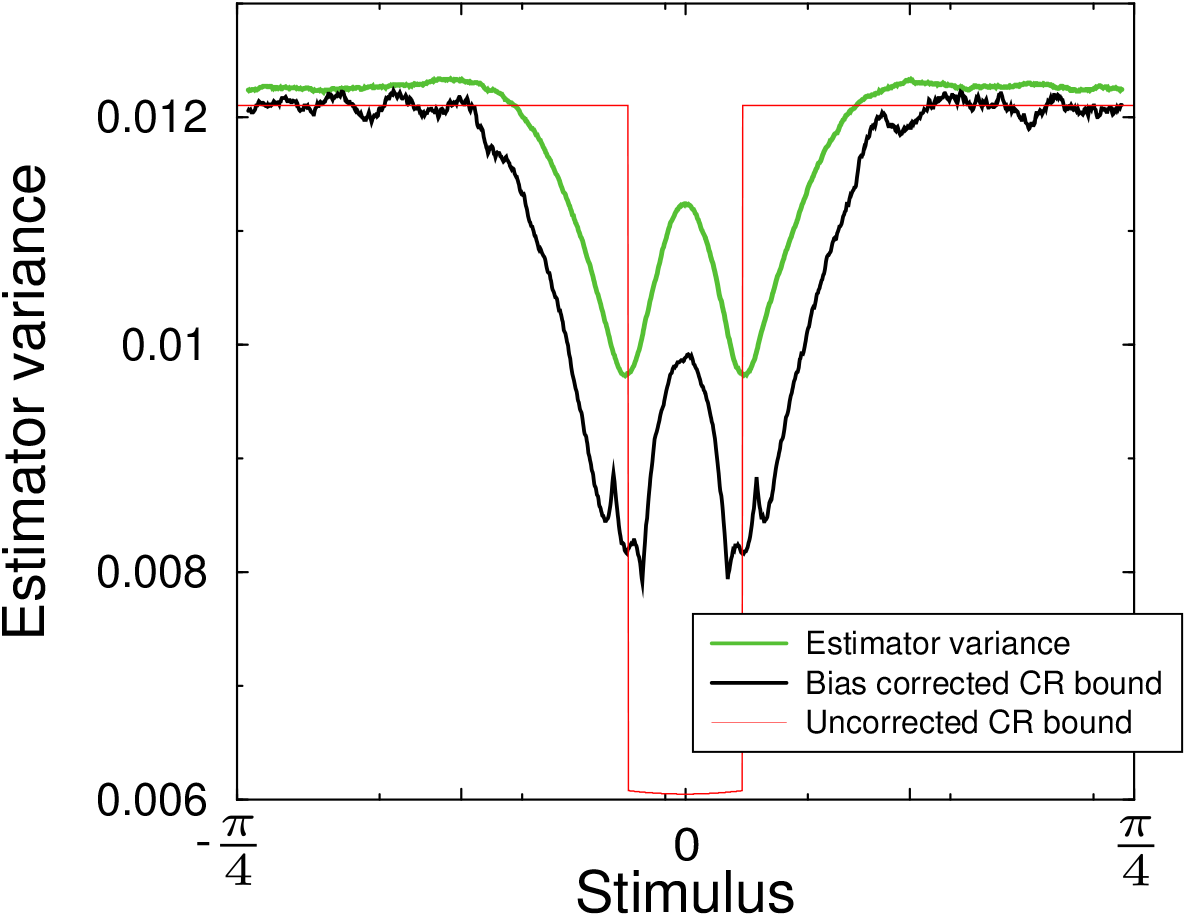
Variance in the Bayesian estimator (green); it is always larger than its lower bound (black). The inverse Fisher information 1*/I*_*F*_ (Θ), which gives the lower bound for an unbiased estimator is also shown (red).

The inverse Fisher information also decreases near the preferred angle (again because the number of active neurons increases). It is essential to incorporate the bias correction in Eq.2, as the variance of the estimator can drop below the inverse Fisher Information (red curve), i.e., break the uncorrected Cramer-Rao bound.

### Approximation for Bayesian estimator bias and variance

Calculating the bias is straightforward, but compute intensive, as it requires to sample many noisy responses (typically 10000 in the figures and even more for larger *N*). For the maximum likelihood estimator we recently introduced a numerically exact way to calculate estimator bias and variance in the case of Gaussian noise. Briefly, one finely discretizes the possible ML candidates and then calculates the probability that a given candidate estimate has the actual maximum likelihood from a high dimensional orthant integral, see Keemink et al. (2018a) for details.

For the Bayesian estimator an decent approximation of the bias can be found as follows. The mean estimate is by definition

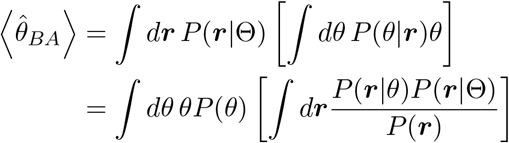

The normalization factor 1*/P* (***r***) prevents doing the integral, but we can approximate the integral by ignoring it. After completing the square, one has 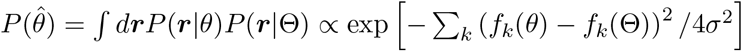.

Assuming a flat prior *P* (*θ*) = *const*., the trial averaged Bayes estimator becomes

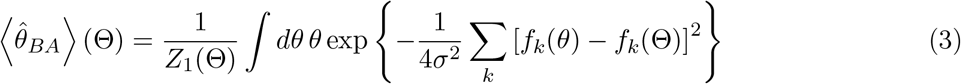

where the normalization 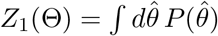.

Similarly, the variance in the estimator is very well approximated with

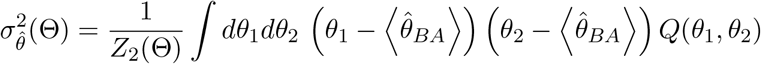

with

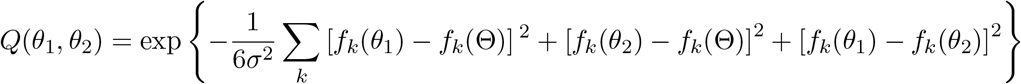

and normalization *Z*_2_(Θ) = *dθ*_1_*dθ*_2_ *Q*(*θ*_1_, *θ*_2_).

These calculations are easily extended to the case of Poisson noise. Here one finds

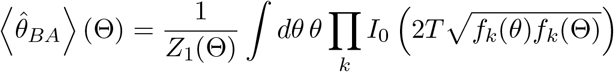

where *I*_0_ is the modified Bessel function of the first kind, *f* is again the tuning curve but now expressed as a firing rate, and *T* is the duration of the spike count window. Likewise

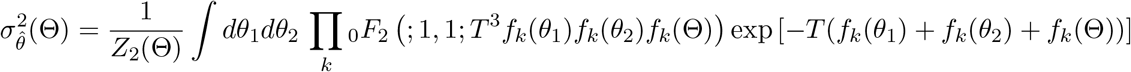

where the generalized hypergeometric function _0_*F*_2_(; 1, 1; *x*) = Σ_*k*_ *x*^*k*^*/*(*k*!)^3^.

To examine the accuracy of this approximation we compare it to the simulation results above, Fig. 6. The approximation follows qualitatively the features observed in the simulation, including its noise dependence and biphasic bias. The approximation however tends to moderately overestimate the bias. Unfortunately, we have no clear insight if or when it breaks down.

**Figure 6:**
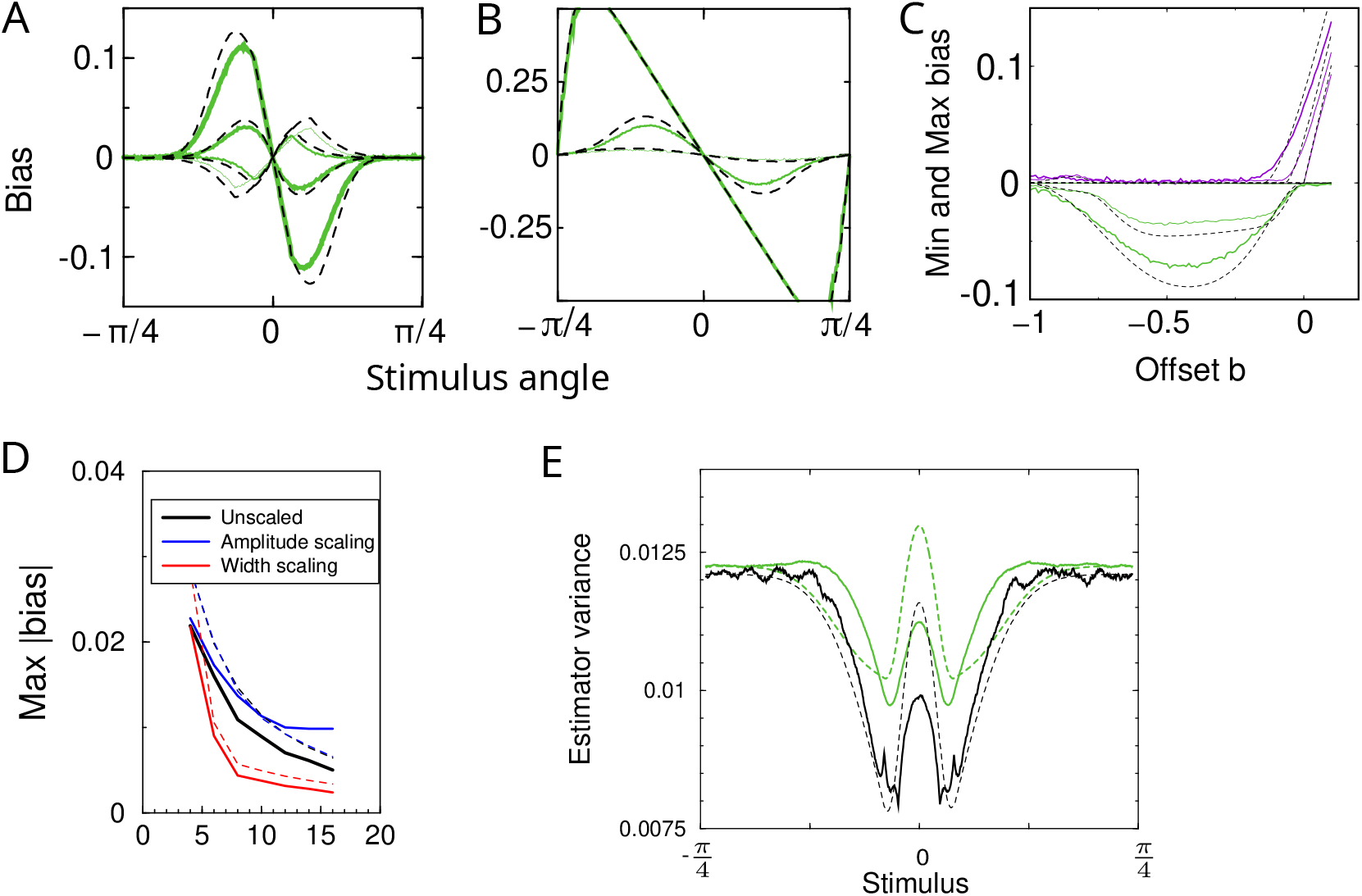
Approximation of the bias and variance of the Bayesian decoder. Simulation results, replotted from above figures are shown as solid lines, the approximation are shown as dashed lines. a) The bias for rectified cos tuning for a number of different offset parameters. b) The bias for von Mises tuning. c) Minimum and maximum bias of cosine rectified tuning for different noise levels. d) Bias against number of neurons. e) Estimator variance and Cramer-Rao bound. All parameters as in previous figures. In panel c) two intermediate noise levels were omitted for clarity.

## Discussion

In summary, population codes that use just a few active neurons are prone to decoding biases. The bias is particularly relevant for system in which just a few neurons are active, i.e. in simple nervous systems found in insects, but also in the retina when very small stimuli are used so that only a few neurons are activated. In particular, the biases of the ML decoder and the Bayesian decoder are similar in size and parameter dependence. But also the population vector shows bias, thus the emergence of bias appears general. Surprisingly, while for von Mises tuning curves the bias is always attractive, for rectified cosine tuning curves the sign of the bias can be either attractive, repulsive, or even bi-phasic. Moreover, while noise always increases bias, the dependence is non-linear and in some cases bias persists at zero noise. The bias is largest for narrow tuning curves. In contrast, it is well known that decoder accuracy increases with narrow tuning (Seung and Sompolinsky, 1993; Zhang and Sejnowski, 1999). The reason is that steep tuning curves yield high sensitivity to small stimulus changes, which reduces variance in the decoder. Thus achieving small decoding bias and small decoding variance might be biologically competing objectives.

The complicated dependence of the bias on tuning curve properties and noise, hinder analytical treatment in even simple cases. To partly mitigate this issue, we have introduced an approximation to the bias and variance in a Bayesian decoder. While this approximation leads to errors in the bias, this does not appear to depend much on parameters. The approximation is efficient as it is just a single integral (per stimulus) and can be carried out with standard integration routines. In contrast, the method in Keemink et al. (2018a) to calculate the bias and variance in a maximum likelihood decoder, is very accurate but still quite computationally demanding as it relies on Monte Carlo integration. The approximation introduced here allows for a rapid calculation of biases, sufficiently accurate for exploratory purposes.

Because the bias is a deterministic function, it is in principle possible to create a decoder that inverts the bias and thus compensates for it. When the bias is constant, compensation is trivial. But here the bias varies with the true stimulus and a compensation mechanism is complicated, in particular because the amount of compensation needs to be aware of noise, i.e. the observation time. If a mismatched bias correction is used, the cure might be worse than the disease. Furthermore, in Keemink et al. (2018a) a Tikhonov regularized bias compensation was employed, which indeed reduced bias, however it led to a large increase in the variance of the decoder. However, in the nervous system, subsequent processing stages do not read out the code, but instead process the population codes. More interesting therefore would be to consider in the future how biases propagate through networks.

Finally we note that biases might also have functional benefits in particular when they are noise dependent. When it is crucial to never underestimate a sensory stimulus, say, in collision avoidance, a noise-dependent bias might lead to an adaptive safety margin. It will be interesting to examine whether such effects are exploited in biology.

## Acknowledgements

It is a pleasure to thank Nikos Gekas for discussion.

